# kontakteUR: transforming coordinates to chemical intuition to focus on essential interactions in biomolecular systems

**DOI:** 10.64898/2026.06.19.732925

**Authors:** Marvin Scherlo, Elias Wippermann, Torben Fürtges, Ricarda Künne, Ayşa Yelboğa, Fabian Rütten, Marcus Böckmann, Udo Höweler, Till Rudack

## Abstract

Molecular interactions govern cellular function, making them essential to discover biomolecular mechanisms by unravelling structure-function relationships. The rapid growth of AI-based prediction, experimental determination, and molecular dynamics simulations generates structural data at an unprecedented scale. However, structural information is typically represented as Cartesian coordinates, leaving chemical interactions and conformational relationships largely implicit. We introduce a high-throughput framework transforming structural geometry into a standardized, compact contact space. Moving beyond simple distance cutoffs, it provides a chemically and geometrically informed representation of various residue–residue interactions, their temporal changes, and conformations at residue-level resolution. Our contact-space representation enables systematic comparison and classification even for large-scale analysis. Case studies spanning structure comparison or studies of protein-protein, protein-ligand, protein-RNA, and antibody–antigen complexes, demonstrate how contact-space analysis reveals interaction patterns, identifies key mutation sites, and links structural features to experimental observations. With these and further applications, kontakteUR elucidates biomolecular function and assists targeted protein design, with results suited for further processing by artificial intelligence algorithms.

## 1 Introduction

The vast majority of processes are driven by a change of molecular contact patterns. Proteins recognize their partners, enzymes position their substrates, signals pass from one domain to the next, and entire molecular machines assemble and disassemble. This holds even for photoreceptors in which a series of light induced structural changes are communicated through interaction networks giving a rise to divers biological function [1]. In large complexes, these interactions are organized into allosteric cycles that synchronize more than one active sites. Examples of such interactions include the stepwise degradation of proteins by the proteasome [2][3] and RNA by the exosome [3][4] or the water splitting process of photosystem II within photosynthesis [5][6].

In most cases the pattern consists of a set of highly specific non-covalent intra– and intermolecular contacts, which form and break during the process. This specificity is explored in protein engineering. Biosensors link a binding event to an optical readout moiety, enzyme and protein design [7] reshape active sites and binding pockets [8][9], and advances in optogenetics and super-resolution microscopy rely on the precise tuning of chromophores [10][11]. Across all these applications, function depends on the specific spacial arrangement of interacting groups and the temporal evolution of the resulting contacts.

Many current applications of interaction analysis rely on simplified distance criteria. Figure 1 shows the results of the prominently used contact definition as any pair of atoms within a fixed cutoff, typically 4 to 6°A between heavy atoms [13][14][15]. This convention is fast and easy to implement, and it remains common in interface analysis and large-scale structural surveys [16][17][18][19][20][21].

**Fig. 1.**
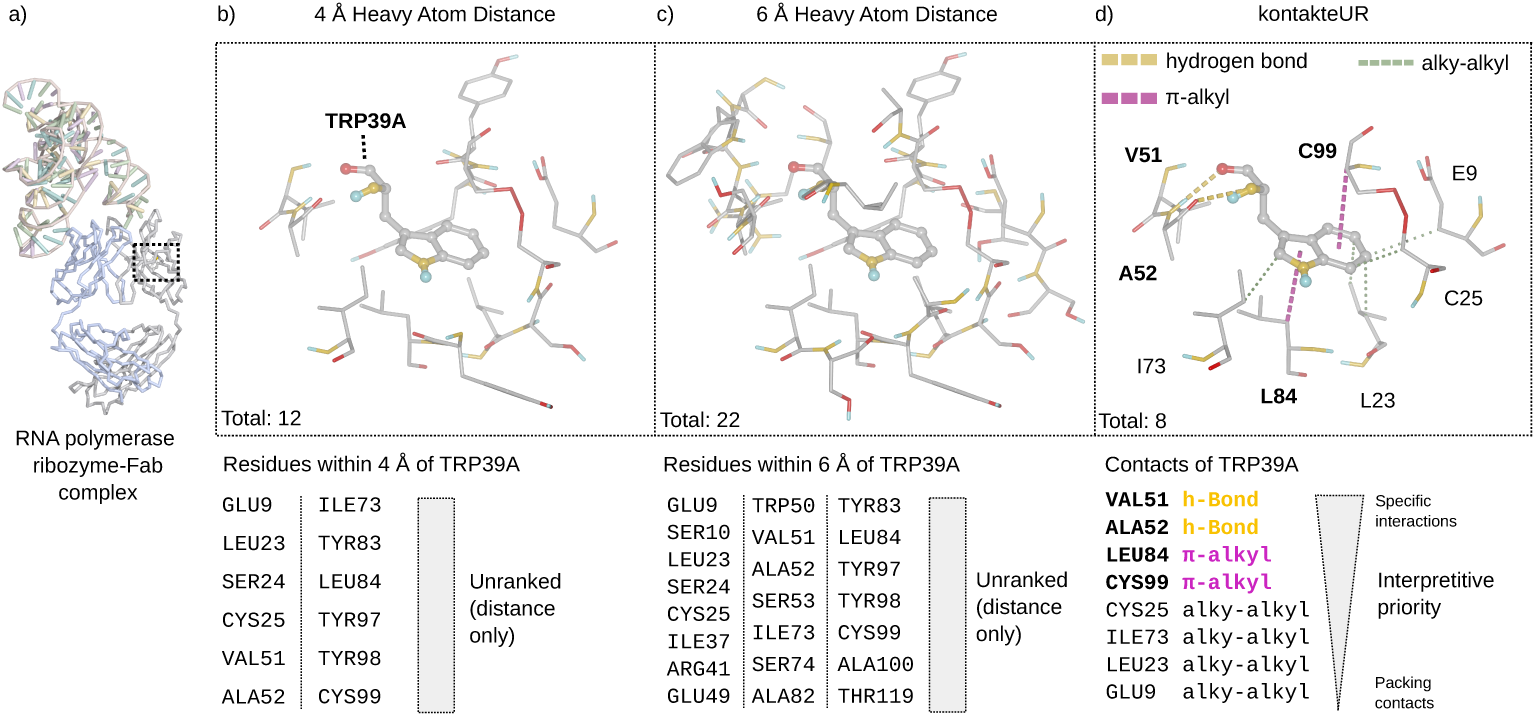
Comparison of distance-based and chemically resolved descriptions of the local environment of Trp39 in chain A of an RNA polymerase ribozyme–antibody Fab complex (PDB ID: 3IVK; [12]). (a) Structural overview. (b, c) Heavy-atom distance cutoffs of 4 and 6°A, representing the lower and upper limits commonly used in the literature, identify 12 and 22 nearby residues, respectively. Because these distance-based analyses neither classify nor prioritize the contacts, key interactions are lost among residues identified solely by spatial proximity. (d) kontakteUR, the contact-analysis tool introduced in this work, identifies eight contact partners and resolves their chemical interaction types, allowing specific interactions to be distinguished from less specific packing contacts. All residue labels refer to chain A.

We report the methodological landscape that has emerged to address the limitations of simple distance criteria and classify these approaches into five functional groups. **I** Static interaction profilers, most prominently PLIP [22] and Arpeggio[23], alongside ligand-focused tools such as LUNA [24] and BINANA 2 [25], employ SMARTS-based atom typing together with interaction-specific geometric criteria to produce chemically detailed annotations of single structures. When applied to molecular dynamics (MD) simulation trajectories, these tools are typically run on a frame by frame basis, treating each frame as an independent structure rather than as part of a coherent time series. **II** The trajectory-native tools GetContacts [26], ProLIF [27], and PyContact [28] operate directly on trajectory formats and generate time-resolved contact information, each defined by a distinct scope: specific system classes, particular interaction types, or contacts between predefined atom selections. **III** Residue interaction network generators such as RING 4.0 [29] and PyInteraph2 [30] aggregate contacts into graphs, abstracting from chemical detail to reveal topological features such as allosteric or signaling pathways. **IV** Visualization tools such as LigPlot+ [31] and iCn3D [32] produce publication-quality two– and three-dimensional representations, prioritizing readability and communication. **V** Energy-based approaches such as INTAA [33] adopt a fundamentally different approach, reformulating contacts as pairwise interaction energies derived from force-field parameters to enable quantitative thermodynamic interpretation.

Each of these tools is well suited to its own purpose. Taken together, they cover much of the relevant methodological space, yet the boundaries between them, and the assumptions each imposes to the data, mean that any specific analysis is shaped as much by the choice of tool as by the underlying question.

We confirm that distances are a necessary but not sufficient criteria to identify a chemically meaningful interaction. For example, two atoms within 5°A may be poorly oriented, shielded by intervening atoms, chemically incompatible, or part of incidental packing. Such arrangements may be tolerated only because favorable interactions elsewhere in the structure outweigh them. More fundamentally, a chemically meaningful interaction cannot, in general, be captured by a single scalar distance. It requires interaction-specific geometric criteria and chemically informed atom types. The practical consequences of replacing interaction-specific criteria with a single distance cutoff are illustrated in Figure 1.

Therefore, the guiding principle of the approach presented here is to preserve a holistic view of the data before deciding which features matter, rather than fixing relevance in advance by restricting the analysis to features already expected to be important. Our principle considers every system component: proteins, nucleic acids, water molecules, ions, lipids, and other hetero groups, including interactions between these classes. It extends across a range of input geometries, including experimentally determined structures, predicted structures, docking poses or frames from trajectories. Filters can then be applied to highlight relevant features, e.g. protein-protein or protein-ligand/drug contacts, which is especially valuable when the meaningful events are not known in advance. We implemented this principle in the here presented tool kontakteUR (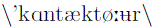).

## 2 Results

The results present a series of versatile case studies illustrating how the application of kantakteUR assists to discover structure-function relationship by extracting meaningful information. First, we introduce the concept of contact space, a change of perspective from atomic coordinates to chemically meaningful interactions and conformational features, followed by the general workflow that enables its efficient generation, filtering, and analysis for structures, ensembles, and molecular dynamics trajectories. Subsequently, we demonstrate its application to diverse biological systems and research questions. Comparison of an experimental and predicted CDK2-CyclinE structure reveals key interchain contacts relevant for structure assessment. Analysis of Ras dimer MD simulation trajectories identifies critical subunit contacts, sporadic and persistent, guiding mutational studies that were later validated experimentally. Investigation of rhodopsin-bound retinal simulation highlights the ability to distinguish different types of *π*-interactions. Examples from enzyme engineering and protein design demonstrate how contact-based analysis uncovers interaction patterns relevant for catalytic activity, substrate recognition, and chromophore spectral tuning. Finally, antibody–antigen interaction analysis illustrates the identification of specific and sporadically occurring contacts from large structural datasets. Such dynamic interaction patterns are also characteristic of many protein–RNA systems, where transient contacts play an important role in molecular recognition.

Together, these diverse case studies show how the contact-space representation draws attention to structural features that are not readily apparent from 3D Cartesian coordinate data alone.

### 2.1 The stringent implementation of the concept of contacts introduces the chemist’s perspective into structure-function relationships

Resolving interactions from a chemist’s perspective requires structural data, and such data is now abundant. X-ray crystallography, cryo-electron microscopy (cryo-EM), artificial intelligence (AI)-driven structure prediction [34][35][36][37], molecular docking approaches, and molecular dynamics (MD) simulations all describe biomolecules as atom positions in three-dimensional space.

This representation captures the geometry fully, but it is not the form in which most biological questions are asked. It describes the system in geometric terms, but not in the functional categories a chemist reasons with. Structure-function relationships are usually discussed in terms of features such as protein or ligand recognition, conformational states, complex formation, and how these change over time. Thus, a chemist’s view requires a transformation from atomic coordinates to functional contacts.

We implement this transformation in our tool kontakteUR, a stand-alone version based on the contact matrix algorithm implemented in the comprehensive molecular modeling package MAXIMOBY [38] and validated through years of practical application. At the core of this implementation is the definition of the contact space. Contacts and conformational features are both be expressed in this space as specific relative arrangements: a contact in how functional groups are positioned toward one another, a conformation in how the atomic centers of a residue are arranged.

The multitude of contacts is organized into chemically distinct interaction types. Atom types are differentiated according to their functionality, allowing contacts to be classified by interacting functional groups rather than by residue identity alone. The resulting matrix records these contact types together with selected conformational features, with backbone and sidechain participation labelled where relevant.

### 2.2 Workflow: Transforming Coordintes into Contacts and Extracting meaningful details

The core functionality of kontakteUR centers on the transformation of structural coordinate data into a contact-based representation and its functional evaluation (Figure 2). Each input structure is converted into a contact vector encoding the pairwise interactions present in that structure together with selected conformational features for each residue. Concatenating these vectors across all input structures produces a comprehensive contact matrix that serves as the primary object of analysis.

**Fig. 2.**
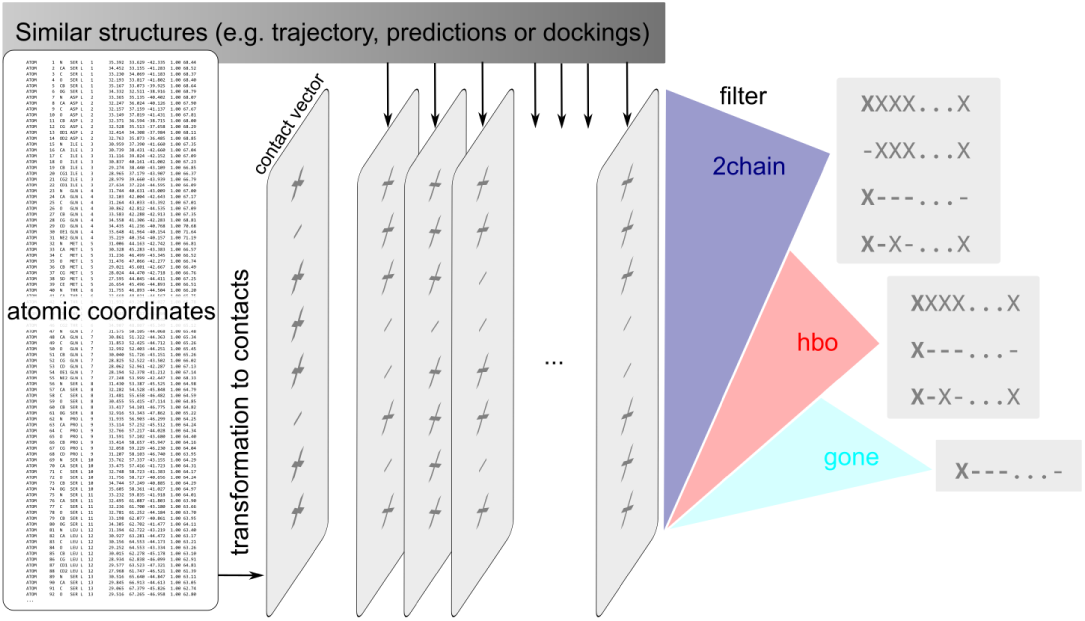
Overview of the kontakteUR workflow. Cartesian coordinates from simulation trajectories or discrete protein conformations are translated into per-structure contact vectors, which are assembled into a contact matrix spanning all input structures. The matrix can subsequently be passed through modular filters to isolate specific interaction types of interest.

**Fig. 3.**
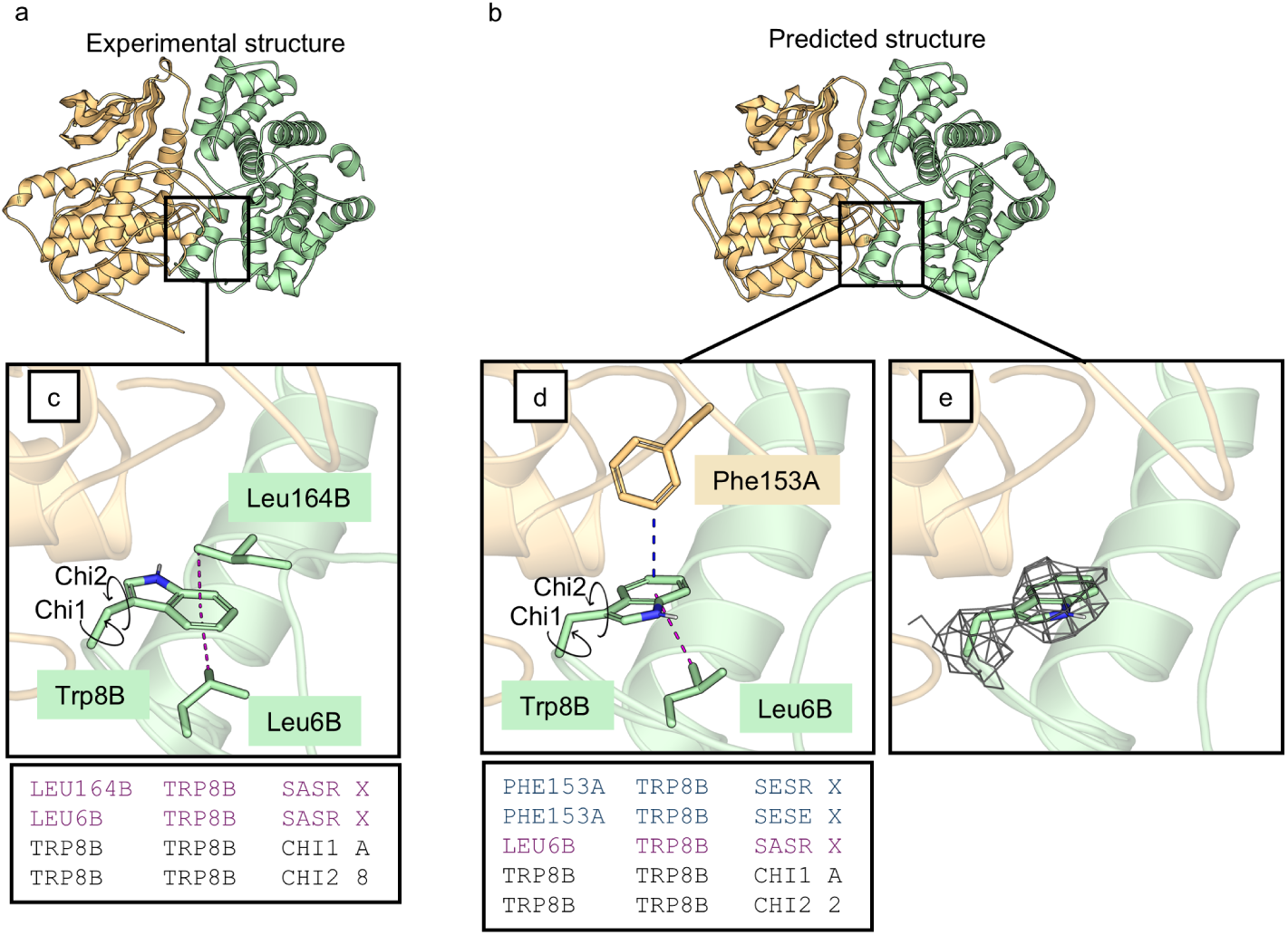
Identifying structural differences by contact and conformational analysis. Comparison of the experimentally derived X-ray structure of the CDK2–CyclinE complex (a) with a corresponding complex predicted by Chai-1 (b). Using our contact analysis metric combined with an interchain filter (2chain), we identified Trp8B in the predicted model as a potential stabilizing residue at the protein–protein interface through interchain *π* ring-ring interactions (d). In contrast, Trp8B in the experimental structure forms only intrachain ring–alkyl contacts (c). Our analysis tool further revealed differences in the side-chain conformation of Trp8B, reflected by distinct Chi2 torsion angles (c,d). Comparison with the experimental electron density showed that the Chai-1-predicted Trp8 side-chain conformation is fully consistent with the observed density (e).

Once contacts and conformational features are gathered in a single matrix, a flexible, modular filtering system highlight those that matter in a given context. Filters can be applied individually or combined in series, using criteria such as persistence, changes between defined temporal or structural subsets, cooccurrence with other contacts and conformational transitions, or participation of selected molecular components. This enables the targeted identification of, for example, hydrogen bonds, ring contacts, inter– or intra-subunit interactions, all contacts involving a ligand, or transient contacts occurring only in a subset of structures. A complete description of the available filters and their parameters is provided in the kontakteUR documentation. The resulting representation is not limited to a particular type of structural data. For trajectories, the contacts of every frame are assembled into a time-resolved matrix, making the temporal development of contacts and conformational features accessible and their correlations visible in a format that is easy to process and interoperable with complementary analyses. A trajectory is only one possible source of such a matrix, however. Any set of structures can be assembled and compared in the same way, including experimentally determined states, models from different prediction methods, or alternative docking poses. Differences and similarities are thus accessible without any simulation at all, and the influence of method or environment becomes directly comparable.

The relevant features therefore emerge from the matrix itself, which is especially valuable when the meaningful events are not known in advance, such as when identifying contacts that stabilize an interface or a ligand, tracing networks of correlated contacts through a structure, comparing predicted, docked, or simulated geometries, or searching for the rare frames in which a transient interaction occurs.

### 2.3 Structure Comparison in Contact Space Reveals Detailed Differences

The example presented here illustrates how transforming structures into contact and conformational space facilitates structure comparison by focusing on chemically meaningful interaction patterns rather than atomic coordinates alone. As shown below, this change in perspective highlights subtle but important differences that are not immediately apparent from visual inspection of the structures.

To compare two structural models of the same protein complex, we analyzed the experimentally determined X-ray structure of the *human* CDK2–cyclin E complex (PDB-ID: 1w98)[39] alongside a corresponding complex predicted by Chai-1 [40]. At first glance, both structures appeared highly similar, with noticeable differences primarily restricted to disordered terminal regions. For a more detailed comparison, both coordinate files were transformed into residue–residue contact representations. This initial step produced extensive contact files listing all non-covalent interactions present within each structure. To focus specifically on protein–protein interface interactions, we subsequently reduced the datasets using a simple text-based filtering command that retained only interchain contacts between CDK2 and CyclinE (2chain fliter, see figure 2).

This analysis highlighted a notable difference involving Trp8 of CyclinE in the predicted structure, where the residue forms an interchain aromatic *π−π* interaction with Phe153 of CDK2. In contrast, no interchain contacts involving Trp8B were detected in the experimental X-ray structure. Instead, the tryptophan side chain adopts an alternative conformation, forming intrachain ring–alkyl interactions with two leucine residues of the same chain. These differences were captured not only by the contact representation but also by the conformational descriptor, which revealed distinct *χ*_2_ torsion angles for the two Trp8 side-chain conformations. Guided by the contact and conformational analysis, we subsequently inspected the corresponding electron density. The side-chain conformation predicted by Chai-1 showed full agreement with the experimental density.

Together, these results demonstrate how transformation into contact and conformational space reduces structural complexity while highlighting chemically meaningful interaction patterns and geometric differences. This facilitates the identification of relevant structural features and guides targeted inspection of residues that may otherwise escape attention during coordinate-based structure comparison.

### 2.4 Embedded Chromophore (Retinal) into Channelrhodopsin II via Specific *π*-System Contacts Over Time

Chromophores, like the retinal, often mediate their biological function through light-induced changes in their conformational states [41][42]. Interactions with the surrounding protein environment can influence chromophore conformation and dynamics, thereby affecting its functional behavior [43]. Analyzing the interactions between the chromophore and surrounding amino acids can therefore help identifying interactions that determine the chromophore’s integration and function within the protein system. A contact analysis of retinal in channelrhodopsin II from Chlamydomonas reinhardtii in the relaxed state reveals, over the course of a one-microsecond MD-simulation [44], a large number of specific-system contacts alongside non-specific van der Waals contacts. These *π*-system contacts play a key role in stabilizing the chromophore (see Figure 4).

**Fig. 4.**
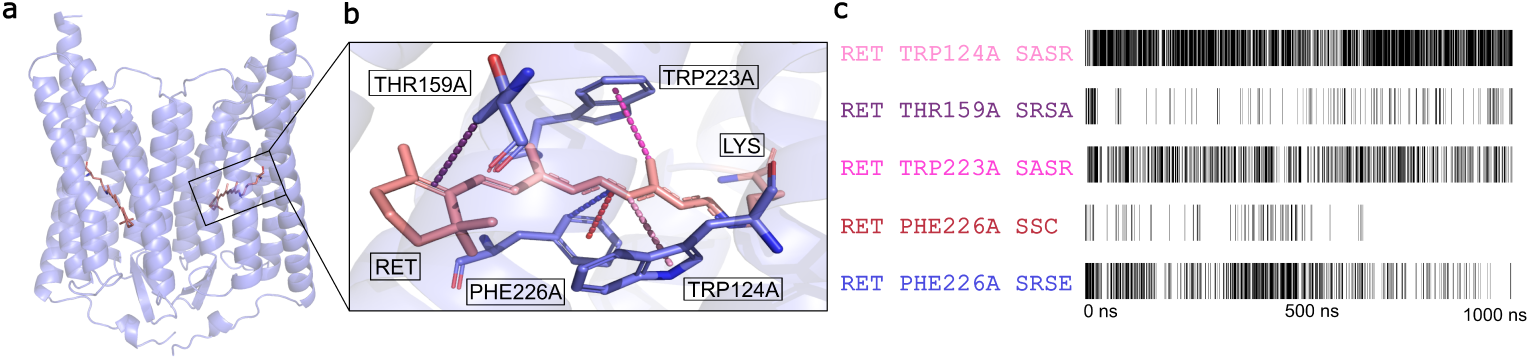
Contact analysis of the retinal in channelrhodopsin II (PDBID:6EID) over time. **a** Overview of channelrhodopsin II containing two retinal molecules covalently bound to the protein. **b** Binding mode of retinal A mediated by specific van der Waals interactions. Retinal is shown in pink, while amino acids involved in contact formation are highlighted in blue. Frequently occurring contacts during the simulation are indicated by dashed lines. **c** Temporal evolution of specific van der Waals contacts during a one-microsecond classical MD simulation. Black lines indicate detected contacts; contacts were evaluated at intervals of 0.5ns.

Notably, the SASR (alkyl-*π*-system, face) contact between retinal and TRP124A remains present for almost the entire duration of the simulation, indicating a highly stable interaction. In contrast, the corresponding interaction between retinal and THR159A occurs only intermittently and therefore appears to contribute less consistently to retinal stabilization. Furthermore, the 6-ring contacts SASR with TRP223A and SRSE with PHE226A exhibit strongly dynamic behaviour, repeatedly forming and dissociating throughout the simulation, suggesting transient stabilization effects. In contrast, the SSC (*π*-system, center– *π*-system, center) contact gradually disappears during the course of the simulation, indicating a loss of structural relevance over time. Characterizing interactions throughout the simulation trajectory enables the identification of key interaction sites, thereby providing a powerful approach to reveal functionally relevant contacts in other retinal proteins and offering potential starting points for the targeted modification of the retinal binding pocket.

### 2.5 Identification of Transient and Persistent Interface Contacts in Ras Dimer Simulations

Molecular dynamics simulations generate vast amounts of coordinate data that can be difficult to interpret directly. This example demonstrates how transformation into contact space reduces this complexity while preserving interaction information, enabling efficient analysis of complete trajectories and the identification of both persistent and transient contacts.

The Ras protein, a central molecular switch in cellular growth signaling whose mutations are frequently associated with tumor development, has been shown to form membrane-anchored dimers, with dimerization being crucial for its function. To investigate this system, we performed multiple 800 ns replica MD simulations of a membrane-bound Ras dimer comprising 234,664 atoms (Figure 5a). The transformation from atomic coordinates to contact space requires approximately 1 s per frame, enabling the analysis of long MD trajectories at meaningful temporal resolution (0.1 ns).

**Fig. 5.**
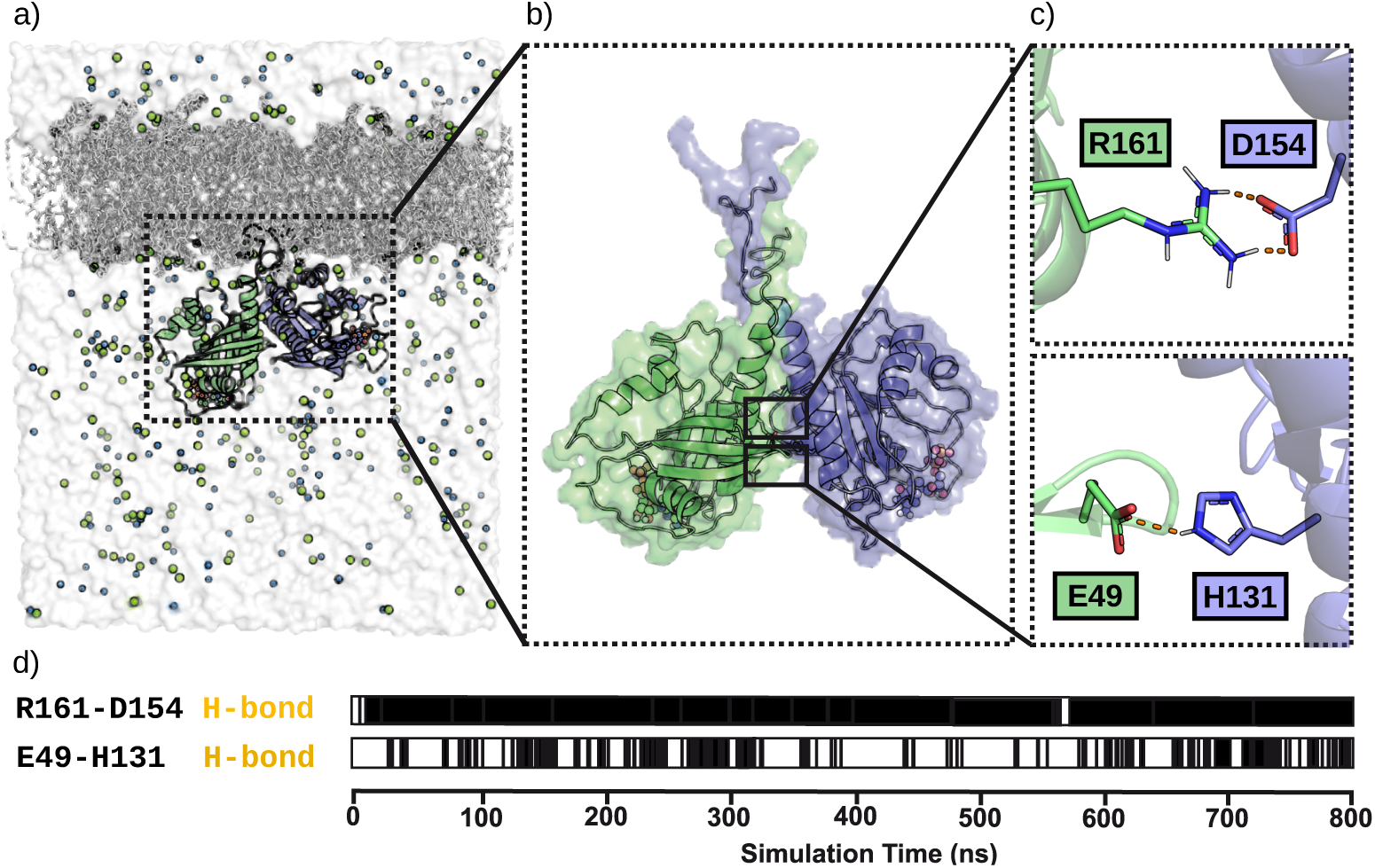
Fast and efficient contact analysis enables the identification of key residues stabilizing the Ras dimer interface from MD simulation trajectories. The simulated system comprises the membrane-bound Ras dimer, lipid bilayer, and solvent, totaling 234,664 atoms (a). Rapid transformation of coordinates into contacts enables analysis of complete trajectories and efficient downstream filtering for interface contacts (b). Subsequent filtering identifies two key hydrogen bonds contributing to dimer stabilization (c). Frame-resolved contact analysis captures both transient and persistent interactions, enabling assessment of contact stability and dynamic behavior throughout the simulation (d). Figure modified after [45].

Combining frame-by-frame contact representations into a contact matrix describing the entire trajectory, we applied the previously described 2chain filter (2.2) to identify residues contributing to stabilization of the dimer interface (Figure 5b-c). The matrix thus identified two key interactions stabilizing the interface. The first, between Glu49 and His131 is transient but persists throughout the simulations and appears relevant for dimer stabilization and shielding of the remaining interface from the water environment due to its position at the interface edge. Such transient interactions are difficult to identify without frame-resolved contact analysis (Figure 5d). The second interaction is a stable hydrogen bond between Arg161 and Asp154 is maintained throughout the simulation and therefore appeared to contribute substantially to dimer stabilization.

These observations enabled the proposal of targeted mutations. Notably, the functional importance of the identified residues was subsequently supported by FRET measurements of the Ras double mutant E49Q/D154N, revealing impaired dimerization [45]. Together, these results demonstrate how contact-space representations transform dynamic coordinate data into interpretable interaction networks, enabling the identification of transient yet functionally important contacts, assessment of their persistence and occurrence frequencies over time, and direct links between molecular simulations and experimental observations.

### 2.6 Investigation of enzymatic activity and protein design

To fully grasp the function of enzymatic proteins a detailed understanding of the molecular mechanism within the active site is crucial. Contacts analysis of active sites have the power to enable such understanding by providing insight into contact pattern across different enzyme-substrate complexes, revealing key residues within reaction intermediates and allow the identification of ambient factors such as water molecules. These findings assist targeted protein design, in which contact analysis again provides a fast and reliable tool to estimate the effect of specific mutations. In the following three case studies contacts analysis was used for the such activity assessment and protein design.

To gain insights into the substrate specificity of proteins of the amidohydrolase superfamily the protein contacts of protein-substrate complexes, intermediate states and protein-product complexes. The analysis elucidates the stabilization of achiral intermediate states, by highlighting a orientation-directing histidin. [46]

Further work addressed the inversion of enantioselectivity in phosphotriesterase through incorporation of a light-responsive unnatural amino acid. The observed loss of contacts between the unnatural aminocads and the protein environment within the predicted substrate-product complexes lead to findings about the mechanism of selectivity inversion. [8]

In a third study the tuning of spectral properties of a biliverdin chromophor through protein mutation was investigated. Multiple variants of bacteriophytochrome with bound biliverdin in *cis* and *trans* state were modeled and evaluated regarding stabilizing contacts. Table 1 depicts shared and differing contacts within the variants. Changes in the types of contacts fitted the observed changes in the spectral properties of the respective variants. [11]

**Table 1.** Excerpt of the contact matrix of biliverdin in across the wildtype (WT) and different protein variants (V1 & V2).

| Contact Partner | Type of Contact | *cis* |  |  | *trans* |  |  |
| --- | --- | --- | --- | --- | --- | --- | --- |
|  |  | WT | V1 | V2 | WT | V1 | V2 |
| His251 | SRSA | - | - | - | - | X | - |
| His251 | SRSE | X | X | X | X | X | X |
| His251 | SSC | - | X | X | - | - | - |

### 2.7 Combined Contact and Conformation Analysis Guides the Design of a Binding-Abolishing Antibody Mutant

Contact analysis also guided the rational design of a binding-abolishing antibody mutant. For the Alzheimer antibody solanezumab bound to its amyloid-*β* antigen, the contact matrix of a crystal structure identified a buried double-phenylalanine motif (F19, F20) as the core of the binding interface (Fig. 6) [47]. Assembling the contacts of several replica simulations into a single time-resolved matrix reduced each trajectory to its essential interactions and made them directly comparable. Across these replicas, only the hydrophobic contacts of the core remained persistent, whereas the surrounding polar contacts fluctuated due to their interaction with the water environment and were absent in some trajectories altogether. The matrix singled out G95^HC^, forming a stable backbone alkyl-*π* contact with F19, as a prominent candidate for mutation to alanine (Figure 6c). The contact matrix further confirmed that the remaining binding pocket was stable across all replicas, leaving little scope for compensatory rearrangements and making even this minimal increase in side-chain size sufficient to disrupt binding.

**Fig. 6.**
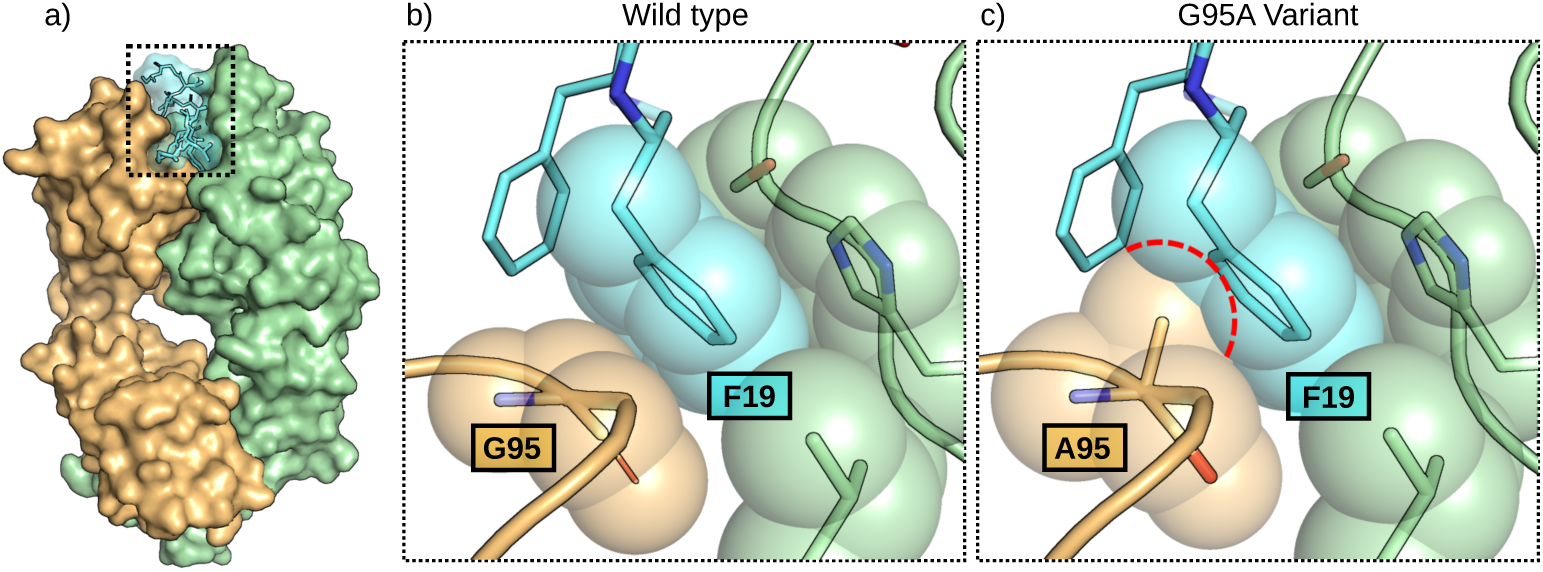
Contact and conformation analysis of the solanezumab–amyloid-*β* interface, the basis for a predicted binding-abolishing mutation. All highlighted interactions and the chosen mutation site were identified from the contact matrix of the complex, computed for the crystal structure (PDB-ID 4XXD) and across replica MD simulations. (A) Solanezumab Fab (heavy chain orange, light chain green) with bound A*β* peptide (cyan). (B) Hydrogen-bond network at the interface (dashed lines), as resolved in the matrix into backbone– and side-chain-specific contacts. (C) The double-phenylalanine motif (F19, F20), which the matrix singled out as the persistent hydrophobic core of the interface: F19 is buried in a pocket formed by G95^HC^ and the light-chain residues H34^LC^, L46^LC^, and S91^LC^. The matrix further reported the backbone torsion of G95^HC^, in the crystal structure and throughout the simulations, as lying within a range that alanine can adopt, identifying it as the most promising pocket-lining residue that could be mutated to abolish without distorting the antibody fold. (D) Predicted G95A^HC^ mutant: the introduced C*β* atom (red dashed outline) overlaps with F19, blocking its insertion and abolishing the binding motif. Figure modified after [47]

The same matrix also confirmed that this glycine could be replaced without structural cost. Because glycine combines the absence of a side chain with unusual backbone flexibility, it is often structurally required in tight loops, where its mutation would distort the local fold. Here, the backbone torsion angles recorded alongside the contacts, both in the crystal structure and throughout the simulations, remained within a range that alanine can equally adopt, showing that the glycine was not conformationally constrained. The mutation could therefore disrupt the binding motif while leaving the loop geometry and the native fold of the antibody intact. The prediction was confirmed experimentally: the mutant lost antigen binding while retaining its native structure.

### 2.8 Summary

The versatility of this approach was demonstrated across a broad range of applications. More importantly, all case studies shared a common outcome: the transformation to contact space highlighted residue-level interaction patterns and conformational features that were directly linked to biological function and experimental observables. For structure comparison, contact-based analysis facilitated the comparison of experimental and predicted structures. In the CyclinE–CDK complex, this perspective enabled the identification of a key interchain contact and a more favorable tryptophan rotamer, providing a clear interpretation of structural differences. Extension of the framework to molecular dynamics trajectories allows for the efficient identification of transient contacts and dynamic interaction networks. Detailed classification of interaction types further enabled the characterization of specific *π*-interactions in retinal-bound rhodopsin that would be difficult to extract systematically from proximity data alone. In the Ras dimer simulations, our approach revealed both stable and sporadically occurring interface contacts that contribute to dimer stabilization, guiding mutational studies that were subsequently confirmed experimentally through the disruption of dimerization.

In general, the residue-level contact representation proved particularly valuable for connecting computational models with experimental observations. In studies of proteins from the amidohydrolase superfamily, contact analysis facilitated the interpretation of protein–substrate interactions associated with enzymatic activity. Similarly, incorporation of a light-responsive unnatural amino acid into phosphotriesterase and the analysis of chromophore–protein interactions in phytochromes demonstrated how contact-based descriptions can relate structural changes to functional and spectroscopic properties, thereby supporting rational protein design of optogenetic tools and biomolecular sensors.

Many biomolecular recognition processes are governed not only by persistent interactions but also by transient, low-lifetime contacts. Such sporadic contacts are of particular relevance for antibody–antigen recognition and are likewise characteristic of many protein–RNA interaction networks, where dynamic contact formation contributes substantially to molecular recognition and function. Our framework enables the analysis of structural ensembles and molecular dynamics trajectories, allowing transient contacts to be captured alongside persistent interactions. A particularly illustrative example was provided by the antibody solanezumab, where contact analysis identified a key interacting residue whose mutation abolished antigen binding. Building on this concept, ongoing work focuses on the analysis of extensive datasets of predicted antibody structures, their dynamics, and docking-derived protein–antigen complexes, where contact patterns are systematically evaluated with respect to specific residues, interaction types, and functional properties.

## 3 Methods

kontakteUR performs a transformation from 3D coordinate-based descriptions of biomolecular systems, like in a pdb file, to a contact scheeme rooted in chemical intuition. Therefore types and forms of contacts as well as a set of conformational properties that complete the description of internal degrees of freedom have to be defined. The methodology described below outlines the chemical principles and implementation of the transformation for each type of information.

### 3.1 Contacts

Independent of the specific type, contacts are generated via a set of spikes that represent an atomic (group) property able to interact favorably with its proximity. The spikes are located at the atoms carrying the property and point into a direction favorable for a position of the complementary property. The tip of the spike marks the *optimal* distance for the pair of properties. With every tip of the spike a catch radius is defined. A contact is detected whenever this radius catches the base point of the complementary spike and vice versa. This principle for detecting contacts via catch radii is illustrated in Figure 7a on the example of a hydrogen bond between the backbones on an alaine (donor) and a phenyl alanine (acceptor). For a hydrogen bond it is essential that the donor atom lies within the catch radius of one of the spikes originating from the acceptor atom **and** the acceptor atom lies within the catch radius of the spike originating from the donor atom.

**Fig. 7.**
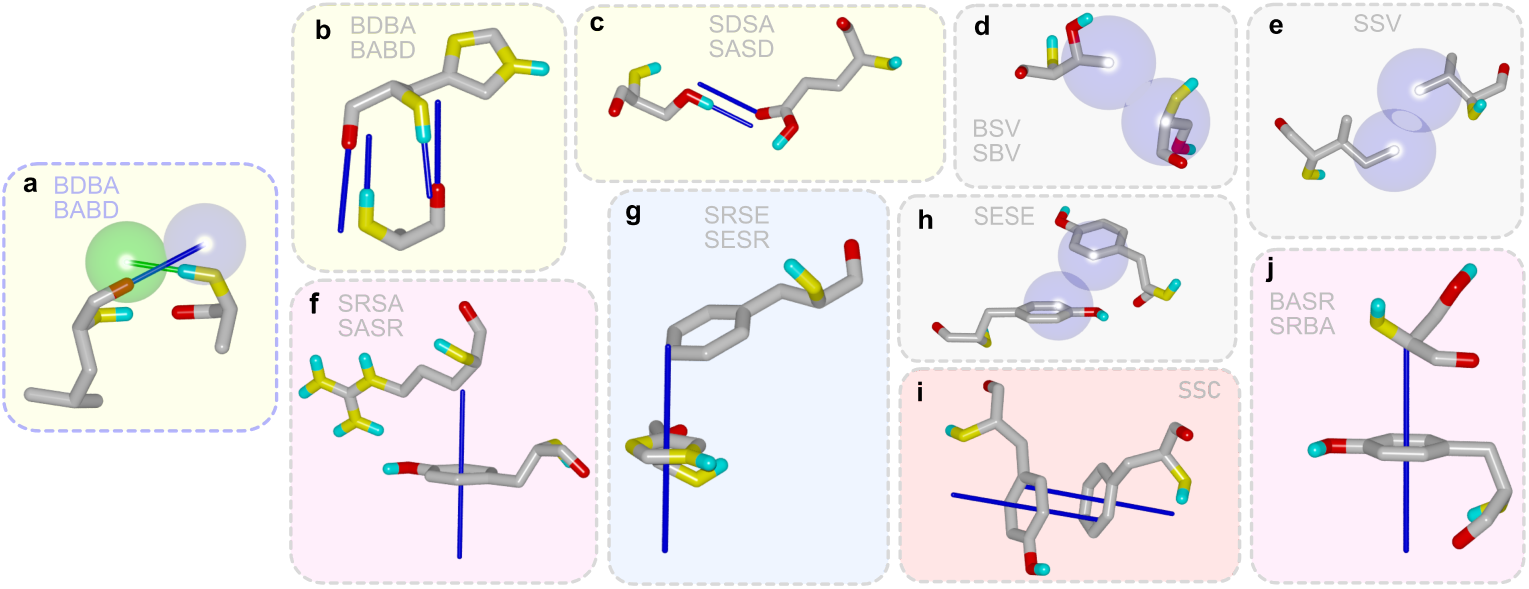
**a** Representation of a hydrogen bond between two backbone atoms, including spikes and catch radii. As the nitrogen lies within the catch radius of one of the five oxygen acceptor spikes (blue) and the oxygen lies within the catch radius of the nitrogen donor spike (green), a hydrogen bond is detected despite less-than-ideal alignment. **b**-**j** Overview of the contact types implemented in kontakteUR. The spikes and van der Waals radii which are causal for the contacts are shown in blue. A description of the individual contacts can be found in the Supplementary Information.

In the SI it is shown for various bond types how wide a spectrum of spike orientations is tolerated to fulfill the combination requirements, demonstrating that rather generously large radii are used to cover all the contacts considered existent.

Following, the different types of contacts are introduced and illustrated in Figure 7. Most contacts are labeled by a four-letter code, where the first two letters represent the first residue and the last two represent the second residue. The first letter in each designation indicates whether the residue belongs to the backbone (B) or the side chain (S). This results in two designations for each contact, depending on the *direction* in which residues one and two are defined. Exceptions are contacts that would be named identically in both directions, such as SSV or SSC. An overview of the various terms can be found in the SI.

#### 3.1.1 Hydrogen Bonds

The donor component of a hydrogen bond is defined by the spike emanating from the heteroatom carrying the hydrogen. The spike passes through the hydrogen atom and has a length that will optimaly find the interacting acceptor. The different type of acceptors carry different sets of spikes *looking* for hydrogen bond donors.

So the oxygen in a R-O-H group carries the donor spike representing the hydrogen and two acceptor spikes at torsion angles (R-1)-R-O-H of +/− 120° with respect to the hydrogen. At the position of an oxygen of the carbonyl-group, which is essential for protein structures, a total of five spikes is used. Two of these spikes point along the idealized positions of the lone pairs in the plane of the R-C(=O)-R (120° to the C=O bond). The next two point at the same angle but perpendicular to the C=O-lone pair plane represent the higher electron density at oxygen due to the *π*-system of the C=O double bond. The set is completed by a fifth spike pointing along the C=O bond simply symbolizing the electron surplus at the oxygen available for hydrogen bonds. Figure 7c (SDSA/SASD) shows a contact between the donor peak of the oxygen in an R-O-H group (serine) and an acceptor peak from a carbonyl group (glutamine acid).

#### 3.1.2 Specific *π*-System Contacts

As directional as the hydrogen bond contacts are the possible interactions with *π*-systems. Usually these *π*-systems represent aromatic rings in protein residues but we extended the scheme to double bonded carbon systems. The *π*-electron coordination for an aromatic ring is implemented by a pair of spikes that emanates from the center of the ring in perpendicular directions to the *π*-system plane. These spikes are used to find favorable interactions with other *π*-systems (stacking), rings (stabilizing edges) and alkyl-groups.

Figure 7f shows such an alkyl contact between the *π*-ring-system of tyrosine and an alkyl in the side chain of an arginine (SRSA/SASR). Similarily, for a *π*-alkyl contact in the backbone, a detected contact between the *π*-ring-system of a tyronine and the *C_α_*atom of a serine is shown in Figure 7j (BASR/SRBA). An exemplary contact to the stabilizing edge of another *π*-system can also be seen in Figure 7 (SES-R/SRSE). The phenylalanine is oriented relative to the histidine so that the corner of the aromatic ring is close to the tip of the spike of the aromatic ring of the histidine. *π*-stacking-arrangements are classified as SCC contacts and illustrated via a tyrosine and a phenylalanine in Figure 7i, stacked on top of each other. One can see that both spikes of the *π*-ring fall on the *π*-ring of the other residue, representing a clear *π*-stacking-arrangement.

Again, the catch radii allow for quite some orientations in these contacts (see Figure A.1 in the SI).

#### 3.1.3 General van der Waals-contacts

Beyond these specifications groups will be in non-directionial contacts. These are defined by very short spikes carrying quite large catch radii representing spherical groups. These can – according to our definition – easily form contacts by simply overlapping and are further differentiated based on the contributing groups of the residue. An example of a van der Waals contact between two side chains (SSV) of a leucin and a valine is shown in Figure 7e. A van der Waals contact between the *C_α_*atom of a serine and a carbon atom in the side chain of threonine is depicted (BSV/SVB) in Figure 7d. When the van der Waals contacts between two *π*-ring edges overlap, as shown in Figure 7h for two tyrosines, this is classified as a separate type of van der Waals contact, namely SESE.

#### 3.1.4 Further structural features

Further *coordinates* in the inverse space comprise the essential properties of amino acid residues and are also identified by kontakteUR (see Figure 8).

**Fig. 8.**
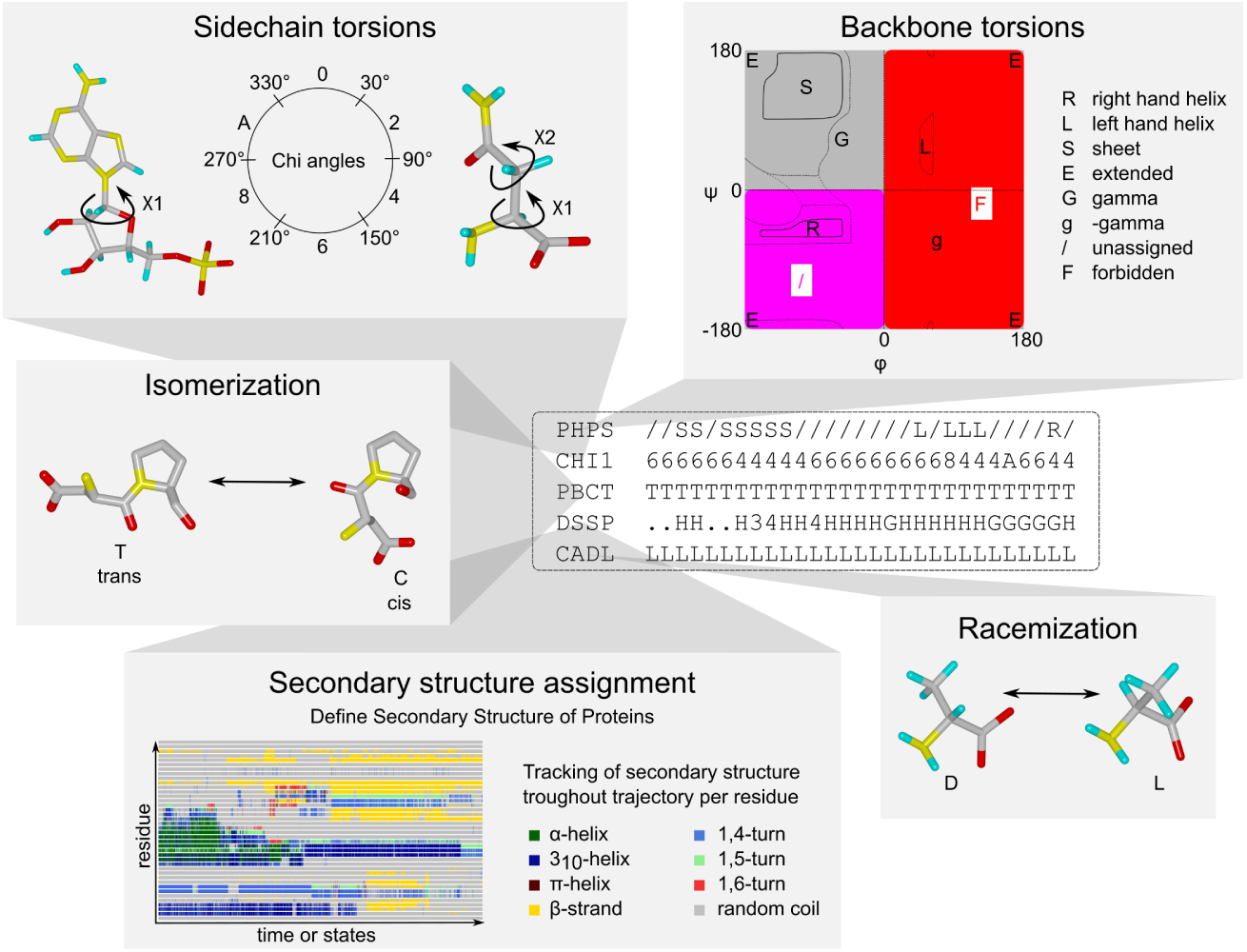
Overview of non-contact structural features encoded in the contact matrix. Residue-level descriptors include secondary structure assignments derived from the Define of Secondary Structure of Proteins (DSSP) algorithm, backbone dihedral angles (*ϕ* and *ψ*), and side-chain rotameric states (*χ* angles). Peptide bonds are characterized through the classification of cis and trans conformations, monitored continuously across trajectories. Stereochemical integrity is assessed via tracking of enantiomeric configurations over time.

The DSSP (Define of Secondary Structure of Proteins)[48] –convention is detected to determine the secondary structure belonging of a residue. By plotting these data on a graph that displays the residues along the input trajectory and color-codes respecting the secondary structure element, it is possible, for example, to directly identify and pinpoint changes in the secondary structure over the course of an MD simulation. In analogy to DSSP-based secondary structure assignment, the Ramachandran convention characterizes the backbone conformation of each residue via its *ϕ* and *ψ* torsion angles. Mapping these angles onto the Ramachandran space allows classification of residue conformations (e.g., *α*-helical, *β*-strand, or coil-like states) and their transitions.

In addition, *χ* side-chain torsion angles are monitored to describe rotameric states of amino acid side chains. An analogous torsional descriptor is defined for nucleic acids, where the torsion angle *χ*_1_ describes the orientation of the nucleobase relative to the sugar moiety. This additional conformational information is required to distinguish Watson–Crick and Hoogsteen adenine–thymine base pairs, which exhibit identical residue-level contact patterns but differ in base orientation (see Figure A3). Finally, stereochemical properties such as the chirality at the *C_α_*atom and the geometry of the peptide bond are assessed to ensure structural consistency, including verification of L-stereochemistry and the predominantly planar trans configuration of peptide bonds.

The various contacts are labeled by a four-letter code specifying the belonging of the interacting groups to the backbone and side chain, respectively. Then all contacts of a single geometry result are gathered in a column called contact vector. By linking multiple structures together in the form of a trajectory (such as time steps in an MD simulation), the contact vector expands into a contact matrix, enabling a direct comparison between structures (see Figure 2).

## 4 Conclusion

We introduce a guiding principle to preserve a holistic view of geometries, a set of geometries, and trajectories by transforming 3D coordinates to chemically relevant contacts. In the basis of this intuitive contact space we validated the applicability of our approach in various biological example systems unraveling structure-function relationships.

Our definition of a contact-based representation of molecular geometry offers a new perspective on molecular structure and function. If the data is presented clearly, it is easier to comprehend. This concept for example allows a human to describe the chromophore binding (Figure 4) as follows: “The retinal binding is dominated by a permanent stabilization by the Trp124 ring, accompanied by methyl group contact to the Trp223 ring while with time the electronically relevant *π*-system stacking with Phe226 switched to a ring edged stabilization. The *π*-system methyl group Thr159 is sporadic but persistent.” without looking at a single geometry.

Nevertheless the vast amount of data in an inter operable format is available for artificial intelligence based algorithmic analysis as it encodes every detail of the geometry.

