## Supplementary material for "kontakteUR: transforming coordinates to chemical intuition to focus on essential interactions in biomolecular systems": Supplemantary Information

**Supplementary Information**

**Table 1** Legend of all differentiated types of contacts.

| Contact Name | Type | Description | Criteria |
|---|---|---|---|
| BDBA | **B**ackbone **D**onor - **B**ackbone **A**cceptor | HBO | Spikes |
| BABD | **B**ackbone **A**cceptor - **B**ackbone **D**onor | HBO | Spikes |
| SDBA | **S**idechain **D**onor - **B**ackbone **A**cceptor | HBO | Spikes |
| SABD | **S**idechain **A**cceptor - **B**ackbone **D**onor | HBO | Spikes |
| BDSA | **B**ackbone **D**onor - **S**idechain **A**cceptor | HBO | Spikes |
| BASD | **B**ackbone **A**cceptor - **S**idechain **D**onor | HBO | Spikes |
| SDSA | **S**idechain **D**onor - **S**idechain **A**cceptor | HBO | Spikes |
| SASD | **S**idechain **A**cceptor - **S**idechain **D**onor | HBO | Spikes |
| SSV | **S**idechain - **S**idechain **V**dW | vdW | Distance |
| BSV | **B**ackbone - **S**idechain **V**dW | vdW | Distance |
| SBV | **B**ackbone - **S**idechain **V**dW | vdW | Distance |
| SESE | **S**idechain ring **E**dge - **S**idechain ring **E**dge | vdW | Distance |
| BASR | **B**ackbone **A**lkylic - **S**idechain **R**ing | $\pi$-contact | Spikes |
| SRBA | **S**idechain **R**ing - **B**ackbone **A**lkylic | $\pi$-contact | Spikes |
| SASR | **S**idechain **A**lkylic - **S**idechain **R**ing | $\pi$-contact | Spikes |
| SRSA | **S**idechain **R**ing - **S**idechain **A**lkylic | $\pi$-contact | Spikes |
| SRSE | **S**idechain **R**ing - **S**idechain ring **E**dge | $\pi$-contact | Spikes |
| SESR | **S**idechain ring **E**dge - **S**idechain **R**ing | $\pi$-contact | Spikes |
| SSC | $\pi$ **S**tacking of **S**ide**C**hain | $\pi$-contact | Spikes |

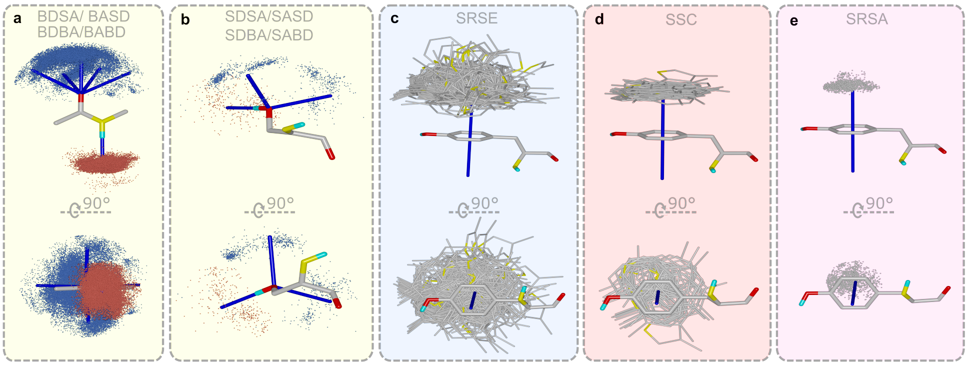


**Fig. 1** Overview of spatial distribution of hydrogen bonds and aromatic ring contacts involving tyrosine across 119 antibody structures. The structures were superimposed by aligned the respective functional group. **a** Detected hydrogen bonds in the backbone. Each point represents the position of an acceptor (red) or donor (blue) partner when the backbone structure is superimposed. **b** Analog representation of hydrogen-bonding partners in an R-O-H group. **c** Ringe-to-Edge (SRSE) contacts, in which the edge of the partner ring is positioned toward the plane of the central tyrosine ring. The corresponding partner rings are displayed in their orientations relatice to the tyrosine ring system. **d** SSC contacts, corresponding to $\pi$-$\pi$-stacking interactions between approximately parallel aromatic ring systems. The corresponding partner rings are displayed in their orientations relatice to the tyrosine ring system. **e** SRSA contacts, corresponding to $\pi$-alkyl contacts between the $\pi$-ring and a sidechain alkyl.

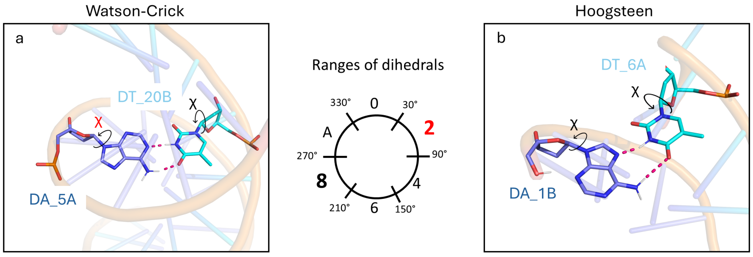


| Residue 1 | Residue 2 | Type |
|---|---|---|
| DA_5A | DT_20B | SASD X |
| DA_5A | DT_20B | SDSA X |
| DT_20B | DT_20B | CHI1 8 |
| **DA_5A** | **DA_5A** | **CHI1 2** |

| Residue 1 | Residue 2 | Type |
|---|---|---|
| DA_1B | DT_6A | SASD X |
| DA_1B | DT_6A | SDSA X |
| DT_6A | DT_6A | CHI1 8 |
| **DA_1B** | **DA_1B** | **CHI1 8** |

**Fig. 2** Combining residue-level contact information and base torsion angles to distinguish DNA base-pairing modes. Watson–Crick base pairing observed in the DNA double-strand X-ray structure PDB 1BNA (a) [**?** ] and Hoogsteen base pairing observed in the X-ray structure PDB 2QS6 (b) [**?** ] both involve two mutual hydrogen bonds between adenine and thymine bases. However, incorporation of the glycosidic base torsion angle $\chi$ enables discrimination between the two pairing geometries due to the distinct adenine rotamer conformations associated with Watson–Crick and Hoogsteen pairing, without requiring additional atomistic-level contact descriptors.
